# Re-calibration of flow cytometry standards for plant genome size estimation

**DOI:** 10.1101/2024.11.11.623134

**Authors:** Abhishek Soni, Robert J Henry

## Abstract

The evolution of long-read sequencing technologies has advanced the development of genome assemblies that are now frequently presented as gap-free or nearly gapless, reflecting substantial improvements in sequencing accuracy and completeness. However, discrepancies between the genome size estimates derived from genome assemblies and flow cytometry create ambiguity regarding the accuracy of these complementary approaches. Accurate genome size estimation via flow cytometry relies on use of an internal standard with a genome of known size. Historically, the genome size of these standards was often calibrated against incomplete genome assemblies or non-plant genomes, such as the human male genome, which was previously considered to be 7 pg but is now known to be around 6.15 pg after 20 years of advancements. Calibrating plant references against non-plant standards is not recommended due to differential staining properties. Therefore, we recalibrated the size of five plant genomes commonly used as reference standards in flow cytometry, by utilizing a recent gapless, telomere-to-telomere (T2T) genome assembly of Nipponbare rice. Our results indicate a significant overestimation of around 20% in previous flow cytometry-based estimates for *Pisum sativum* and *Nicotiana benthamiana*, around 10% for *Arabidopsis thaliana*, and less than 5% for *Sorghum bicolor*, and *Gossypium hirsutum*. The close alignment of the recalibrated GS estimates to the reference genome assemblies and recalculated estimates from different studies confirms their suitability as reference standards for more accurate measurement of plant genome size.

## Introduction

Flow cytometry (FCM) is a standard approach for the estimation of plant genome size (GS) by measuring the relative fluorescence of fluorochrome-labelled nucleus against an internal standard of known GS (Galbraith et al. 1983; Doležel et al. 2007; Sliwinska et al. 2022; Temsch et al. 2022; Soni et al. 2024). The GS is typically expressed in C-values, where the DNA content of a non-replicated haploid nucleus is referred to as the “1C-value.” This value can be measured in picograms or million base pairs, with one picogram being equivalent to 978 million base pairs (Doležel et al. 2003). The accuracy of GS estimates is critical in genome sequencing, evolutionary studies, breeding, and biodiversity conservation for efficient allocation of sequencing resources and understanding of the adaptive significance of GS variations.

The accuracy of FCM-based GS estimation depends on selecting internal standards with precisely and accurately known GS values (Bennett and Smith 1976; Doležel and Bartoš 2005; Doležel and Greilhuber 2010; Sliwinska et al. 2022; Temsch et al. 2022). Traditionally, the GS of internal standards was verified by comparing the relative fluorescence of various standards, often including reference genomes of non-plant organisms such as human and chicken (Van’T Hof 1965; Bennett and Smith 1976; Bennett and Smith 1991; Doležel et al. 1998; Praça-Fontes et al. 2011). However, an accurate estimate of the GS of the reference standards requires the achievement of complete genome assemblies (Doležel and Greilhuber 2010) which have only recently become possible (Nurk et al. 2022; Chen et al. 2023; Gladman et al. 2023; Shang et al. 2023; Mo et al. 2024). In addition, the calibration of a plant standard against non-plant genomes may not be reliable due to differential staining properties (Doležel and Greilhuber 2010; Sliwinska et al. 2022) suggesting the use of plants as primary standards to calibrate the GS of plant reference standards. Here, we used *Oryza sativa* ssp. Japonica cv. Nipponbare as a primary standard, given the availability of a complete genome assembly (Shang et al. 2023; Abdullah et al. 2024), to recalibrate five plant standards with ∼27 fold diversity of GS ranging from 0.288 to 7.65 pg. Our estimates were in accordance with the reference genome assemblies of the respective species and showed up to ∼19% difference from previous GS estimates. Previous calibration of several standards against the human male genome (using a size of 2C = 7 pg) (Tiersch et al. 1989), compared to the updated value of 6.15 pg (Nurk et al. 2022), contributed to the earlier overestimations in the reference standards.

## Results

### Recalibrated GS estimates

Using the rice genome assembly as a primary standard, the recalibrated 2C value for *Arabidopasis thaliana*, 0.288±0.002 pg, was approximately 3% smaller than the previous estimate (2C = 0.297 pg) based on chicken erythrocytes (2C = 2.33 pg) (Table 1). The recent genome assemblies with 2C = 0.291, 0.286 pg supported the precision and accuracy of the recalibrated values (Table 1). The recalibrated GS of sorghum (2C = 1.545 pg) based on rice, was nearly 4% larger than the previous, recalculated and T2T genome assembly-based estimates (Table 1). Based on the recalibrated sorghum, cotton was calibrated at 4.733 pg/2C which was about 1.4% smaller than the previous FCM based GS estimate based on chicken erythrocytes (2C = 2.33 pg). Although, the recalibrated value aligned well with the genome assembly-based estimate (2C = 4.75pg) (Table 1). Using recalibrated cotton, the 2C value of pea (7.65 pg) was found to be approximately 15.8% smaller than the previous estimate of 9.09 pg, which was based on the previous human male GS estimate (2C = 7 pg). Moreover, the recalibrated estimate was 19.3% smaller than the previously reported FCM based estimate for the specific Torstag cultivar (2C =9.49 pg) (Jordan et al. 2015). The recalculated GS estimate for pea (2C =7.99pg) against updated human male GS (2C = 6.15 pg) (Doležel et al. 1998), was about 4.2% larger than the recalibrated estimate. In addition, genome assemblies for two other cultivars of pea (2C = 7.67, 7.61pg) corresponded well with the recalibrated as well as recalculated estimate (Table 1). Using updated pea GS, the recalibrated 2C estimate of *Nicotiana benthamiana,* was 5.828pg, approximately 10% smaller than the previous Feulgen densitometry-based estimate (2C = 6.4 pg), which was calibrated against onion (2C = 33.5pg). Although the recalibrated value corresponded well with the recent T2T genome assemblies (2C = 5.826 pg) (Table 1).

**Table 1:**
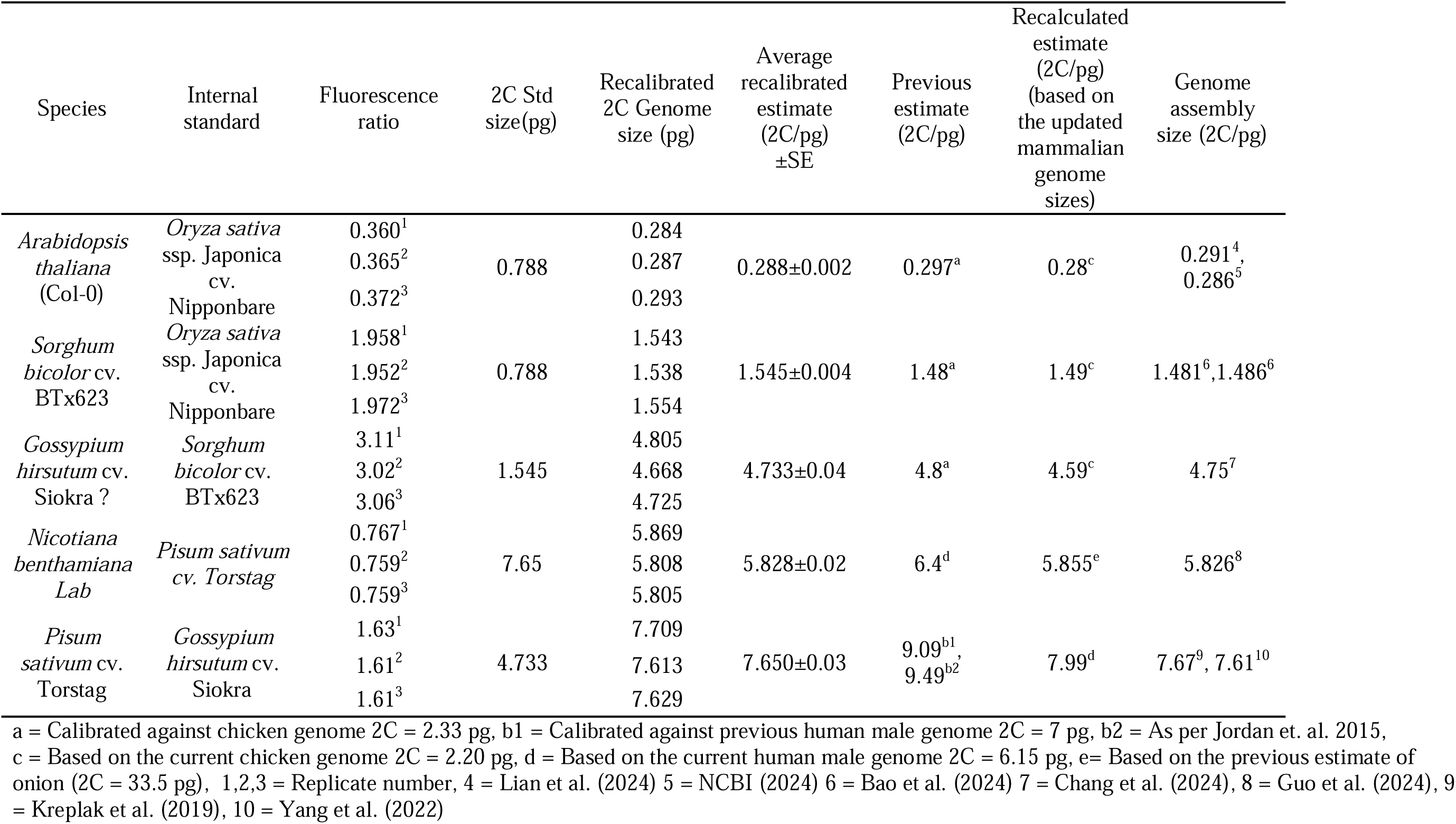
The recalibrated GS estimates, previous estimates, recalculated estimates and corresponding genome assembly sizes.

Regression analysis (Fig 1, 1S) revealed a strong linear relationship between the recalibrated estimates and the other calibration methods, with adjusted R² values ranging from 0.98 to 1 indicating consistency between recalibrated estimates, genome assembly size and the recalculated estimates based on updated mammalian genome sizes, suggesting near-equivalence between these methods. However, the slopes indicated that previous GS estimates were overestimated when compared to the recalibrated estimates (Fig 1). Although, the genome assembly-based estimates and recalculated estimates were marginally different from the recalibrated estimates (Fig 1).

**Figure 1.**
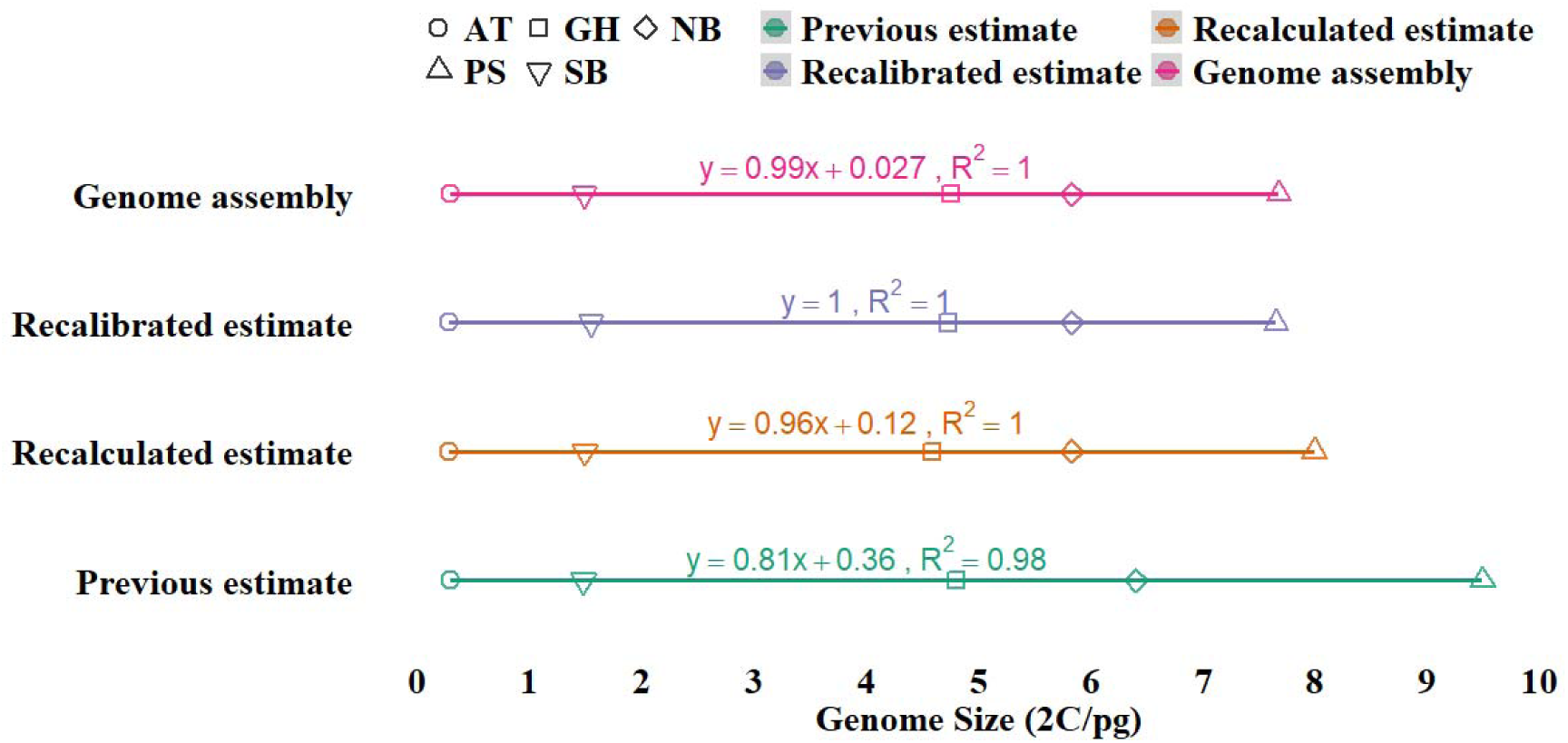
Comparative analysis of recalibrated estimate, recalculated estimate, and genome assembly-based estimate. The graph shows the genome size (2C/pg) of five plant species in four categories: GS from the most complete genome assembly (pink), previous estimate calibrated against human or chicken genome using FCM (green), recalibrated estimate derived by FCM against rice as primary standard (purple), recalculated estimate evaluated by the previous fluorescence ratio and updated GS value of human and chicken genomes (orange). Symbols represent specific plant species: *Arabidopsis thaliana* (Circle, AT), *Sorghum bicolor* (Inverted Triangle, SB), *Gossypium hirsutum* (Square, GH), *Nicotiana benthamiana* (Diamond, NB), *Pisum sativum* (Triangle, PS). The lines represent linear regressions with equations and R² values.

### Discrepancies in the previous GS estimates

Previously, methods like Feulgen Microdensitometry, Spectrophotometry, and cell cycle analysis contributed to the foundational work in GS estimation and the calibration of reference standards (Van’T Hof 1965; Bennett and Smith 1976). Feulgen Microdensitometry involved staining the DNA and quantifying the dye bound to the nuclei, while spectrophotometry measured light absorbance to estimate DNA amounts, and cell cycle analysis inferred GS based on the timing of replication and division phases. However, the accuracy of these methods relied heavily on the correct GS of the reference standard. For instance, Van’T Hof (1965) estimated the GS for onion (2C = 33.5 pg) based on the length of the cell cycle and proportionality of the frequency of 2C, intermediate and 4C cells to the respective duration of the cell cycle stages. However, this method was later found to be less accurate to estimate GS in absolute values (Bennett and Smith 1976). Bennett and Smith (1976) used onion (2C = 33.5 pg) to calibrate eight angiosperm species to be nominated as internal standards, including pea (2C = 9.72 pg). Later, GS of pea varied three-folds in several studies employing different GS estimation techniques (Fig 2) (Lyndon 1963; Bennett and Smith 1976; Bennett and Smith 1991; Michaelson et al. 1991; Doležel et al. 1992; Baranyi et al. 1996; Doležel et al. 1998; Johnston et al. 1999; Doležel and Greilhuber 2010; Kreplak et al. 2019; Yang et al. 2022). However, due to the lack of quality plant genomes, these values could not be verified for accuracy. Although, given their precision from different studies, the values were often opted to estimate the GS of plant species.

**Figure 2.**
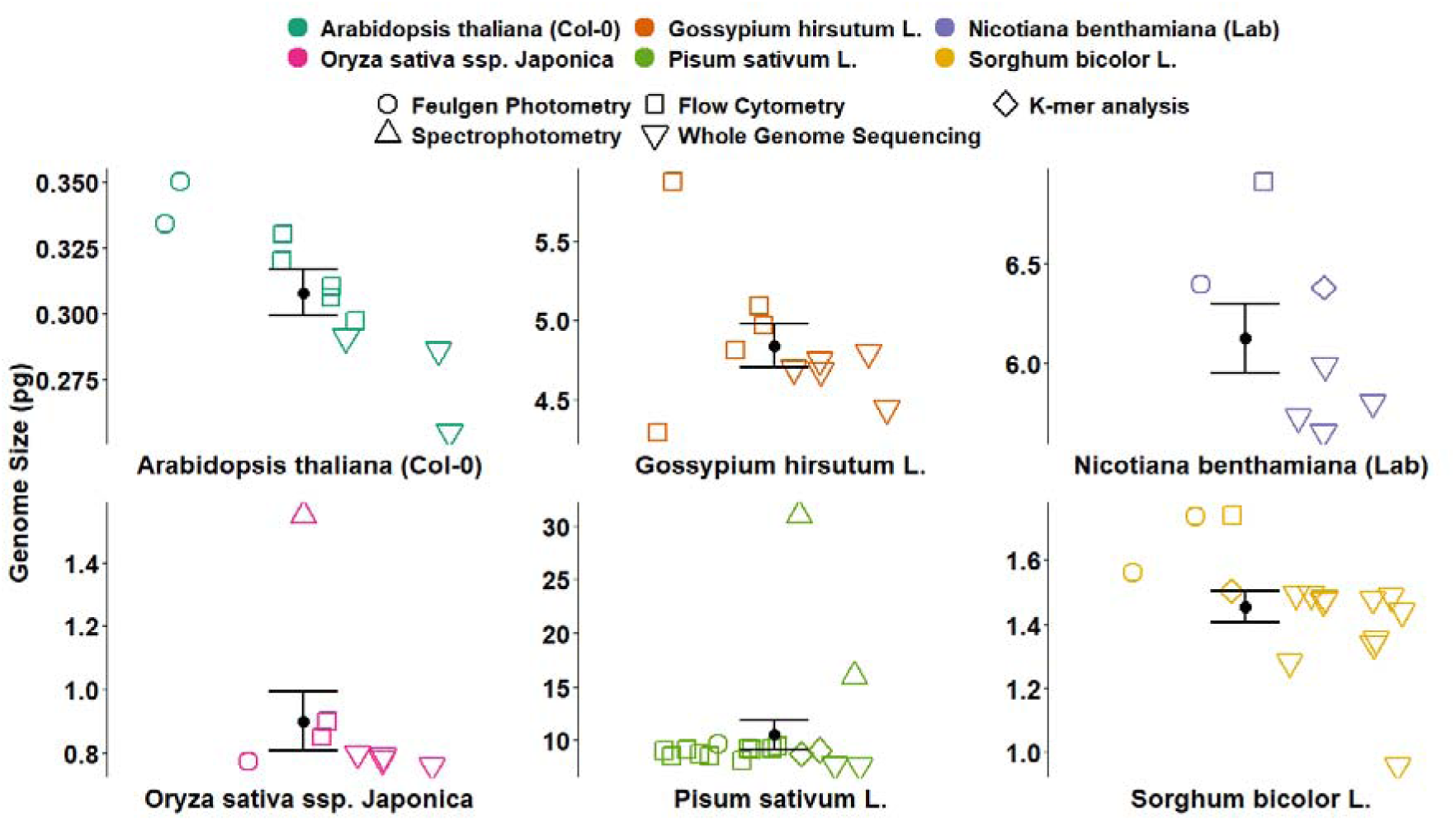
The variability in the genome size (2C/pg) of six plant reference standards derived using several methods in the past.

The calibration of pea against human male leukocytes (2C = 7 pg) as reported by Tiersch et al. (1989) demonstrated high precision, leading to the establishment of pea (2C = 9.09 pg) as a reference standard (Doležel et al. 1992; Doležel et al. 1998). This value has since been widely applied across a variety of plant species for GS estimation and as a guide for genome sequencing studies of pea (Kreplak et al. 2019; Yang et al. 2022).

Similarly, discrepancies from two to three-fold can be observed in the calibrated GS of the common plant standards (Fig 2). The GS of *Arabidopsis thaliana* in previous studies has varied from 0.255-0.334 pg (Bennett and Smith 1991; Galbraith et al. 1991; Krisai and Greilhuber 1997; The Arabidopsis Genome 2000; Bennett 2003; Hou et al. 2022; Kang et al. 2023; Lian et al. 2024; NCBI 2024), allotetraploid cotton varies from 4.29-5.09 pg (Arumuganathan and Earle 1991; Hendrix and Stewart 2005; Li et al. 2015; Hu et al. 2019; Wang et al. 2019; Huang et al. 2020; Chang et al. 2024), *Nicotiana benthamiana* LAB varies from 5.64-6.92 pg (Narayan 1987; Hussain et al. 2021; Kurotani et al. 2023; Ranawaka et al. 2023; Guo et al. 2024; Ko et al. 2024; Wang et al. 2024), sorghum varies from 0.975-1.735 pg (Laurie and Bennett 1985; Johnston et al. 1999; Paterson et al. 2009; Deschamps et al. 2018; Mccormick et al. 2018; Tao et al. 2021; Wang et al. 2021; Bao et al. 2024; Ding et al. 2024), and *O. sativa* ssp. Japonica cv. Nipponbare varies from 0.77-1.55 pg (Bennett and Smith 1976; Iyengar and Sen 1978; Arumuganathan and Earle 1991; Bennett and Smith 1991; Goff et al. 2002; Sasaki 2005; Shang et al. 2023; Abdullah et al. 2024) (Fig 2). Based on the recalibrated GS estimates, the discrepancies in past GS estimates can be attributed to the inaccurate calibration of reference standards, such as the outdated human male genome size of 2C = 7 pg or the chicken genome size of 2C = 2.33 pg, as well as the lack of verification against plant genomes. The recalibration based on the updated mammalian GS and the complete rice genome highlights the extent to which prior GS estimates may have been overestimated.

### The progress in the human genome sequencing

The advent of new sequencing technologies has facilitated the resolution of the complex regions of human genome including centromeric satellite repeats, transposable elements, complex structural variation, Chromosome Y and X (including long palindromes, tandem repeats and segmental duplications (Lander et al. 2001; Schneider et al. 2017; Aganezov et al. 2022; Altemose et al. 2022; Morales et al. 2022; Nurk et al. 2022; Vollger et al. 2022; Chao et al. 2023a; Rhie et al. 2023; Li and Durbin 2024) (Table 2). One of the recently published human genomes, T2T-CHM13v2.0, is now considered gapless and telomere-to-telomere (T2T) (Aganezov et al. 2022; Nurk et al. 2022). With more recent advances and fully sequenced X and Y chromosomes, the current gapless human male genome (44+XY) is around 2C = 6.15 pg (Miga et al. 2020; Nurk et al. 2022; Rhie et al. 2023). Chao et al. (2023b) successfully achieved a gapless genome assembly of a Chinese individual, which closely aligned with the genome size derived from the T2T-CHM13 reference genome assembly. Jarvis et al. (2022) achieved several diploid and haplotype resolved assemblies using a range of sequencing platforms and assembling approaches including CLR reads, CCS reads, ONT reads, ONT Ultra Long reads, Hi-C reads and Bio Nano optimal maps with an assembly size ranging from 5.72 to 6.34 pg in accordance with the expected 2C genome of male 6.16 pg (Table 2). These values were in accordance with the size of the reference genome (T2T-CHM13 v2.0) (Nurk et al., 2022). This high level of congruence demonstrates the precision of genome assembly in representing the human genome. Furthermore, the genome assembly (HG002 Ref.pat) from Jarvis et al. (2022) is ∼100% complete when compared to the T2T-CHM13. Likewise, one of the earlier genomes using Sanger sequencing (GRh38.p13) was also 99.53% of the T2T-CHM13 genome (Jarvis et al., 2022). Assembly completeness of the T2T-CHM13 was assessed using k-mer statistics and mapping-based statistics (Nurk et al., 2022). A comprehensive analysis revealed that the T2T-CHM13 assembly maintains uniform coverage across the genome, with 99.86% of the sequence within three standard deviations of the mean coverage for HiFi and ONT reads (Nurk et al. 2022). Excluding the ribosomal DNA (rDNA) sequences, this uniformity further improves to 99.99%. Despite this high level of completeness, some regions of the genome remain associated with potential issues due to low coverage, low confidence, or known heterozygous sites due to the limitations of both ONT and HiFi read challenges in dealing with telomeric regions and GC rich regions respectively (Nurk et al. 2022; Li and Durbin 2024). These potential issues encompass only 0.3% of the total assembly length, equivalent to approximately 9.165 Mbp, a marked improvement from the 8% of problematic regions observed in GRCh38 (Nurk et al. 2022). This ensures that future research and applications can be guided with greater accuracy and confidence, setting a new standard for genome assembly completeness. Both Han1 (2C = 6.12 pg) and T2T.CHM13v.2.0 (2C = 6.15 pg) are the most complete human genome assemblies (Nurk et al. 2022; Chao et al. 2023a). Considering the reference assemblies and potential sequencing errors in T2T.CHM13v2.0, the complete human male genome was assumed to be 6.15 pg/2C (Table 2). This value agrees with the previous speculations (Doležel et al. 2003; Dolezel et al. 2007). Based on this value GS estimates for plant species were recalculated (Table 2).

**Table 2:** The genome size of a recently assembled human male (44Autosomes+XY) genome.

| Assembly name | 2C value (pg) | Sequencing platform | Reference |
| --- | --- | --- | --- |
| Han1 | 6.12 | PacBio high-fidelity (HiFi) reads, and Oxford Nanopore Technology (ONT) reads. | (Chao et al. 2023a) |
| T2T-CHM13 v.2.0 | 6.15 | PacBio HiFi CCS reads, PacBio CLR reads, Oxford Nanopore Ultra Long Reads, Illumina short reads, 10x Genomics reads, Hi-C Arima Genomics, Bio Nano optical Maps, Single-Cell DNA temple strand sequencing | (Miga et al. 2020; Nurk et al. 2022; Rhie et al. 2023) |
| HG002 Ref.pat | 6.16 | PacBio HiFi, PacBio CLR, ONT Ultra Long 100Kb+, 10x linked reads, Strand-seq, Hi-C linked reads, Illumina, Bio Nano optical maps and Strand-seq) | (Jarvis et al. 2022) |
| GRCH38.p14 | 6.10 | Shotgun sequencing | (Schneider et al. 2017; Morales et al. 2022) |
| GRCH38 | 5.96 | Shotgun sequencing | (Lander et al. 2001) |

### Effect of the updated mammalian genome on the previous GS estimates

Tiersch et al. (1989), calibrated several reference standards against the human male genome (2C = 7 pg) including, chicken genome (2C = 2.33 pg). Both human and chicken genomes have been extensively used as primary standards for verifying the GS for plant internal standards (Bachmann 1972; Galbraith et al. 1983; Tiersch et al. 1989; Doležel et al. 1998; Doležel and Greilhuber 2010) (Table 3). Considering the current human genome size (2C = 6.15 pg) and ratio of chicken/human ratio of 0.357 from Tiersch et al. (1989), the size of chicken should be 2.20 pg. All, current available genome assemblies of chicken highly close to 2.20 pg (references from NCBI). According to the current human genome size, the previous GS of chicken genome (2.33pg) (Galbraith et al. 1983; Tiersch et al. 1989; Arumuganathan and Earle 1991; Doležel and Greilhuber 2010) would be overestimated by ∼6%. Therefore, the recalibrated values for the previously calibrated standard could vary accordingly (Table 3).

**Table 3.**
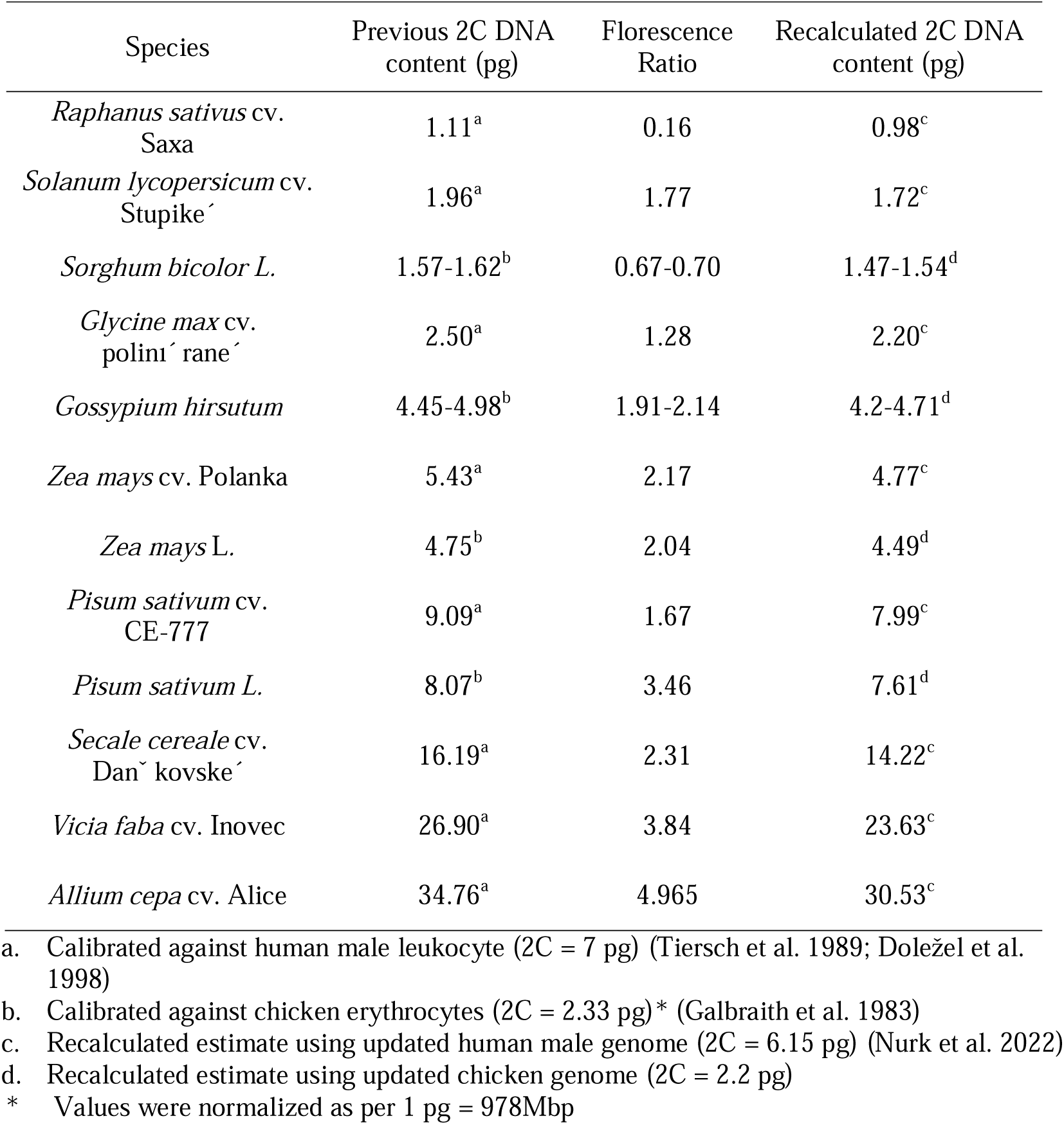
Difference of the previous and recalculated DNA content of the plant reference standards calibrated against old and current genome estimates of human male leukocyte. and chicken erythrocyte.

## Discussion

Flow cytometry is commonly used in genome sequencing studies for resource management and quality checks of assemblies, but its reliability depends heavily on the accuracy of the reference standards. Here we present evidence that inaccuracies in reference standards, such as the overestimation of the human male GS (previously 2C = 7 pg, now corrected to 2C = 6.15 pg), have led to miscalibrations, affecting values for other standards like chicken and plant standards, in particular, pea (Tiersch et al. 1989; Doležel and Bartoš 2005; Doležel and Greilhuber 2010).

In this study, we proposed new calibration values for five plant standards using the complete telomere-to-telomere (T2T) genome of Nipponbare rice, due to its complete and accurate assembly, multiple precise genome assemblies, and suitability as a plant standard with similar GC content (Temsch et al. 2022). Similar to the human genome, the genome of Nipponbare rice was assembled using a hybrid assembly strategy including PacBio HiFi, ultra-long Oxford Nanopore Technology (ONT), and Hi-C reads (Shang et al. 2023). High mapping rates > 99.6% of raw reads to the assembled genome, the presence of 99.88% of the BUSCO gene set and telomere to telomere (T2T) chromosomes represent the completeness of the Nipponbare rice genome (Shang et al. 2023). Some gaps in the Nipponbare genome were previously filled with the gap-free genome ZS97RS3 of *Oryza sativa* spp. Indica genome (Song et al. 2021) and complex repetitive regions such as 45S rDNA arrays were resolved as per the methods for complete human and maize genome assemblies (Nurk et al. 2022; Chen et al. 2023) The estimates of rice genome size have varied from 0.77-1.55 pg in previous studies (Iyengar and Sen 1978; Arumuganathan and Earle 1991; Bennett and Smith 1991; Goff et al. 2002; Sasaki 2005; Shang et al. 2023; Abdullah et al. 2024). Arumuganathan and Earle (1991) estimated 2C = 0.86 pg as lower limit of *O. sativa* against chicken erythrocyte (2C = 2.33 pg). Considering the chicken genome (∼2.2 pg), the rice genome would be 0.800 pg, aligning more closely with the complete genome assembly (Arumuganathan and Earle 1991; Shang et al. 2023; Abdullah et al. 2024).

Based on the Arabidopsis/chicken ratio from Arumuganathan and Earle (1991) and the updated chicken GS (2C = 2.2 pg), the recalculated GS estimate of *A. thaliana* (2C = 0.28 pg) closely matches the recalibrated GS in this study. Similarly, using the Arabidopsis/pea ratio from lamp-based FCM from Doležel et al. (1998) and the recalibrated pea GS, the recalculated estimate of 0.286 pg for *A. thaliana* aligns closely with the recalibrated value. Recent complete T2T genome assembly Col-CC (GCA_028009825.2) NCBI (2024); Reiser et al. (2024) showing 2C values of 0.291 pg and 2C/0.286 pg Lian et al. (2024) further validate the recalibrated values. Krisai and Greilhuber (1997) calibrated size of *A. thaliana* (2C =0. 334 pg) against pea (2C = 8.84 pg) using Feulgen photometry, however considering the recalibrated GS of Pea from this study (2C = 7. 65 pg), the GS of *A. thaliana* would be 0.289 pg/2C. Moreover, 2C value of 0.31 pg, originally calibrated against chicken (2C = 2.33 pg) by Galbraith et al. (1991), adjusts to 0.292 pg when using the revised chicken genome value of 2.2 pg. This adjusted value too aligns closely with the 0.288 pg reported in our study. Our estimates was slightly lower than previous figures of 2C = 0.306 pg against *Drosophila melanogaster, 2C = 0.32 against Caenorhabditis elegans* and 2C = 0.33 pg against chicken erythrocytes (Bennett 2003). Recalculations using updated values for the chicken genome suggest a closer alignment with plant-based calibrations, highlighting the importance of using species-specific standards for accuracy. Given that *A. thaliana* differs from non-plant species significantly in genomic composition, including repetitive DNA and centromeric content, using rice, a fellow angiosperm, provides a more suitable basis for calibration.

The genome size of sorghum varied from 1.28-1.56 pg historically (Laurie and Bennett 1985; Johnston et al. 1999; Paterson et al. 2009; Deschamps et al. 2018; Mccormick et al. 2018; Tao et al. 2021; Wang et al. 2021; Bao et al. 2024; Ding et al. 2024). Although in this study the GS value of 1.545 pg differed ∼14% and ∼ 18% from the genome assemblies, 2C/1.339 pg and 2C/1.279 pg, for BTx623 cultivar assembled with short reads and sanger sequencing respectively (Paterson et al. 2009; Mccormick et al. 2018). Although recent T2T genome assemblies (∼1.49 pg) of sorghum with long read technologies were in accordance with the recalibrated GS value (Bao et al. 2024; Ding et al. 2024). Current T2T assemblies were mapped against raw reads for the completeness and accuracy evaluation. Moreover, the current assemblies with a base error rate ∼1bp per 10Mb are significantly improved from the error rate of 1bp/100kb in earlier genomes (Bao et al. 2024).

The GS of allotetraploid cotton in this study was calibrated at 4.73 pg against sorghum (2C = 1.545 pg). This value was about 5% and 24% smaller than the previous calibration against *O. sativa* IR36 (2C = 1.01 pg) and *Hordeum vulgare* cv. Sultan (2C = 11.12 pg) respectively (Hendrix and Stewart 2005). However, recent genome assemblies using long read sequencing technologies citing 2C = 4.75 pg from Chang et al. (2024), 2C = 4.68 pg from Huang et al. (2020), 2C = 4.80 pg from Wang et al. (2019) and 2C = 4.694 pg from Hu et al. (2019) align close the recalibrated GS.

The recalibrated GS for Torstag cultivar of pea was 7.65 pg/2C, which is about 24% smaller than the earlier reported value of 9.49 pg for the same cultivar used in GS estimation of the Proteaceae family (Jordan et al. 2015). Using the updated human male GS (6.15 pg), the GS value for pea, based on the pea/human ratio of (Doležel et al. 1998), should be 7.99 pg which shows a nearly 14% overestimation in the previous calibration value of pea. Moreover, the recent genome assemblies of pea (2C = 7.61,7.67 pg) reflect accordance with the recalibrated value. Arumuganathan and Earle (1991) reported a genome size of 8.07 pg for pea using a chicken genome size of 2.33 pg, which would be adjusted to 7.63 pg using the chicken genome size of 2.2 pg. Similarly, Johnston et al. (1999) calibrated pea (2C = 9.21 pg) using chicken erythrocytes (2C = 3.01 pg) however, this 2C value of pea would be approximately 7.13 pg with the updated chicken genome.

Lastly, *Nicotiana benthamiana* Lab accession was calibrated at 5.828 pg which matched closely with the recent high quality T2T, complete genome assembly (2C = 5.826 pg) from (Guo et al. 2024). In addition, genomes assembled with long read technologies citing 5.648 pg from Ko et al. (2024), 5.988 pg from Wang et al. (2024), 5.797 pg from Ranawaka et al. (2023) and 5.724 pg from Kurotani et al. (2023) were also in accordance with the recalibrated estimate.

The accuracy of the recalibrated estimates can be verified from the previous results from different labs and methodologies. For instance, the GS of onion was estimated as 35.76 pg/2C and 37.13pg/2C against pea (2C = 9.09 pg) in two laboratories using lamp based FCM (Doležel et al. 1998). When adjusted with the recalibrated pea (2C = 7.65 pg), these estimates reduce to 30.1 pg and 31.2 pg respectively, (averaging 30.65pg). This value closely aligns with Feulgen densitometry based measurements of 33.5 ± 10 % pg by Bennett and Smith (1976). Moreover, the 2C value of onion calibrated using the human male genome (previously 2C = 7 pg) was 34.76 pg. However, when recalculated with the updated human GS (6.15 pg), the recalculated estimate is reduced to 30.54 pg, closely aligning with results from different laboratories and GS measurement techniques (Bennett and Smith 1976; Doležel et al. 1992; Doležel et al. 1998). Likewise, the close resemblance observed between the recalibrated estimate based on laser based FCM (0.288 pg) and recalculated GS estimates (0.286 pg) of *A. thaliana* from different labs (using recalibrate pea at 7.65 pg), as determined by Doležel et al. (1998) using lamp-based FCM. These results eliminate concerns about potential variations between lamp and laser-based flow cytometers (Doležel et al. 1998). The recalculated estimate for pea against onion (2C = 30.65 pg) yields values of 7.79 pg and 7.5 pg from two different labs (Doležel et al. 1998), closely aligning with the recalibrated GS estimate of 7.65 pg. Michaelson et al. (1991) calibrated *Hordeum vulgare* (2C = 11.12 pg) against onion (2C = 33.5 pg). Using the recalculated estimate of 30.65 pg for onion, the GS of *H. vulgare* should be 10.17 pg (Bennett and Smith 1976; Michaelson et al. 1991). This adjustment reduces pea (7.82 pg) and sorghum (1.59 pg) which are in accordance with the recalibrated estimates, thus, verifying the accuracy of recalibrated values against decades-old results from various labs and instruments (Michaelson et al. 1991; Doležel et al. 1998). Likewise, the recalculated estimate for *Z. mays* 4.66 pg and 4.48 pg using recalibrated pea size and fluorescence ratios from Doležel et al. (1998) will be in accordance with the recalculated estimate (2C = 4.49 pg) against updated human genome (Doležel et al. 1992) and a complete genome assembly based estimate (4.45pg) (Chen et al. 2023). Similarly, the recalculated estimate for *Vicia faba* (2C = 23.63 pg) against the updated human genome (Doležel et al. 1992) closely aligns with recalibrated values from Doležel et al. (1998). *Vicia faba* was estimated at 26 pg/2C and 25.19 pg/2C against onion (33.5 pg), and 27.75 pg and 27.92 pg against pea (9.09 pg) from four different laboratories (Doležel et al. 1998). Considering the updated GS values for onion (30.65 pg) and pea (7.65 pg), the recalibrated estimates of 23.78 pg and 23.05 pg against onion, and 23.35 pg and 23.5 pg against pea, are closely aligned with the recalculated estimate of 23.63 pg.

The overestimation of GS, particularly in species like pea, could inflate resource allocation during genome sequencing projects. For instance, the critically endangered *Eidothea hardeniana* was previously estimated 1.81 pg/2C based on the overestimated GS of pea (2C = 9.49 pg) (Jordan et al. 2015; Leitch IJ 2019). However, based on rice (2C = 0.788 pg), we estimated the GS of *Eidothea* to be 1.31 pg, which is 37% smaller than the previous estimate (data not presented here). A telomere-to-telomere (T2T) assembly of *Eidothea* (1.26 pg) aligns closely with current FCM derived estimate (data not presented here).

Finally, the precision of the genome estimates based on FCM, and genome assemblies demonstrate the progress made at the sequencing level and the suitability of genome assemblies for recalibration of flow cytometry standards. However, the consideration of extra-chromosomal DNA (eccDNA) should not be overlooked, as genome assemblies might be representing only the linear chromosomal DNA. In rice, approximately 25,600 eccDNA molecules were identified in various tissues, with nearly 87% found in leaves (Cuzzoni et al. 1990; Zhuang et al. 2024). The length of eccDNA ranged from 74 bp – 5000 bp, with the majority between 200-400 bp. Assuming an average length of 300 bp and approximate 22,180 eccDNA molecules from leaves, would account for 6.65 Mbp. Cuzzoni et al. (1990) estimated eccDNA to be representing 1% of the total DNA. Similarly, the presence of eccDNA has been reported in *Arabidopsis* and *Nicotiana* also (Kinoshita et al. 1985). It will be interesting how flow cytometry (FCM) measurements and genome assemblies consider eccDNA. Given the diverse nature of genome assemblies and eccDNA, genome assemblies might be smaller than FCM estimates due to the exclusion of eccDNA from linear DNA. However, the extent of this difference can only be quantified once eccDNA is identified for each species.

## Methods

### Selection of species

Young seedlings of *Oryza sativa ssp.* Japonica cv. Nipponbare, *Sorghum bicolor cv. BTx623*, and *Pisum sativum* cv. Torstag were cultivated under glasshouse conditions at the University of Queensland, while *Arabidopsis thaliana Col-0*, N*icotiana benthamiana Lab*, and *Gossypium hirsutum* cv. Siokra were grown at the Queensland University of Technology.

We opted these species given their easy accessibility, availability of previously reported GS estimates through different equipment, techniques and laboratories, high-quality genomes assemblies of long-read sequencing technologies and their extensive previous or potential use as reference standards for GS estimation of plants using flow cytometry. Given that the fluorescence ratio between sample/standard should not be more than ± 3-times and should not be overlapping (Dolezel et al. 2007; Sliwinska et al. 2022). To avoid the linearity in fluorescence signal detection, poor resolution (Doležel et al. 1998), we used species that differed by approximately 20-30% in GS from the standard. We employed a sequential calibration strategy with rice as the primary standard we calibrated sorghum and *A*. *thaliana*. Recalibrated sorghum was used to estimate the GS of cotton, and subsequently, recalibrated cotton was used to recalibrate pea which was later used to estimate GS of *Nicotiana benthamiana*.

### Selection of primary standard

Nipponbare rice, with a complete and telomere-to-telomere (T2T) genome (2C = 0.788 pg) Shang et al. (2023), was used as the primary reference standard to calibrate the GS of the other species. The rice genome, assembled using a hybrid approach of HiFi reads for high accuracy and ultra-long ONT reads for resolving complex, repetitive regions, achieved a consensus accuracy of approximately one error per 5 million bases (Q63) (Zhang et al. 2022; Sahu and Liu 2023; Shang et al. 2023; Xie et al. 2024). Similarly, hybrid sequencing approaches have been employed to assemble gapless *Oryza* genomes (Song et al. 2021; Zhang et al. 2022). In addition, Abdullah et al. (2024) assembled a version of the Nipponbare genome with one missing telomere, the GS estimate (2C = 0.781 pg) was in accordance with the complete genome, confirming the accuracy of this estimate.

### Flow cytometry

A one-step protocol was used to isolate intact nuclei from fresh plant material (Galbraith et al. 1983; Doležel et al. 2007; Soni et al. 2024). About 40 mg of both the internal standard and test species were co-chopped in 800 µL of ice-cold modified buffer (Mb01) which was prepared as per (Sadhu et al. 2016). The nuclei suspension was filtered through a 40 µm nylon cell filter and 400 µL of the suspension was transferred to a 5 mL round-bottom FACS tube. A staining buffer consisting of 20 µL Propidium Iodide (1mg/mL) (∼50 ppm final concentration) and 0.2 µL RNase (1mg/mL) was added to the suspension, and the tube was kept on ice until fluorescence measurement. Propidium iodide-labelled nuclei were excited with a 488 nm blue laser and fluorescence was recorded using a Becton Dickinson LSR Fortessa X20 Cell Analyzer equipped with a 695/50 nm bandpass filter (Koutecký et al. 2023). Data were collected on a linear scale with a trigger threshold set at 5000, recording at least 600 events per peak and a total of 2000 events at a flow rate of 12 µL/min, yielding 10-20 events per second. Signal amplification was achieved by setting FSC detector voltage to 320, SSC voltage to 179, and fluorescence detector voltage to 405. For Arabidopsis, the latter was set to 495. Forward and side scatter were recorded on a logarithmic scale to identify fluorescence peaks, and pulse width vs. pulse height plots were used to eliminate aggregates and debris. Consistent manual gating was applied to remove debris across replicates, with a focus on selecting the minimum required number of events from the middle of the population distribution and coefficient of variation (CV) less than 5% (Loureiro et al. 2007). To ensure the accuracy of recalibration, three biological replicates per species were performed to calculate the average genome size. Genome size was calculated as per equation (1). Subsequently, the recalibrated estimates were cross validated against the genome assembly data, previous estimates and recalculated estimates using update human genome size.

Genome size (2C/ pg) = (Mean fluorescence of sample species/Mean fluorescence of internal standard) x 2C (pg) value of internal standard ….1

### Comparison of genome size estimates

To investigate the variability in genome size (GS) estimates of the reference standards, a comprehensive literature review was conducted. The GS estimates derived using various methods including FCM, genome sequencing, k-mer analysis, spectrophotometry and Feulgen microdensitometry were extracted from literature. In addition, C-value database (https://cvalues.science.kew.org/) was used to retrieve previous GS estimates (Leitch IJ 2019) and the NCBI database (https://www.ncbi.nlm.nih.gov/home/genomes/) was searched for the most complete genome assemblies of the selected species. The GS values were standardised and converted to pg/2C values using the conversion factor of 1 pg = 978 Mbp (Doležel et al. 2003). Subsequently, the diversity in these estimates was visualized using R (Ihaka and Gentleman 1996; Wickham 2011). The historical use of the human male genome was explored for calibration of the plant reference standards. Data for the reference human genomes (T2T-CHM13 v.2.0, HG002 Ref.pat, GRCH38.p14, GRCH38, Han1) was retrieved from NCBI genome database. The quality of the genome assemblies and the sequencing technologies were assessed through literature review. Based on the updated human GS (2C = 6.15 pg), the GS of chicken was re-estimated according to the chicken/human ratio provided by Tiersch et al. (1989), resulting in a recalculated 2C estimate of 2.2 pg for chicken. Using the updated values for the human male genome (2C = 6.15 pg) and chicken genome (2C = 2.2 pg), along with fluorescence ratios of plant standards calibrated against human or chicken genomes, the previous GS estimates were recalculated. These recalculated estimates served as a guide for our FCM measurements, allowing for a comparison between previous estimates and the recalibrated estimates from this study. Finally, the recalibrated GS estimates were compared with previous GS estimates, recalculated GS estimates, and the GS estimates from the most complete genome assemblies using linear regression analysis. Since in previous calibration studies, GS of the reference standard was calibrated against different species (Doležel et al. 1998). To satisfy this criterion, we used the pervious fluorescence data from the literature conducted on different machines and against different standards to validate the accuracy of the recalibrated values.

## Data access

The data analysed in this manuscript is presented in the figures, tables, and supplementary materials. The flow cytometry experiment files are available in the “FlowRepository.org” database under the accession number FR-FCM-Z8BJ, and can be accessed directly via the link: http://flowrepository.org/id/FR-FCM-Z8BJ.

## Acknowledgements

We would like to sincerely acknowledge Dr Buddhini Ranawaka for kindly growing the plants of *N. benthamiana, A. thaliana, G. hirsutum* and providing the fresh leaves for flow cytometry. We also thank Prof. Peter Waterhouse for his valuable suggestions for selection of the species for recalibration.

## Author contributions

*AS*: Conceptualization, methodology, writing - original draft preparation, review & editing.

*RH*: Conceptualization, project supervision, funding acquisition, writing - review & editing.

## References

Abdullah M, Furtado A, Masouleh AK, Okemo P, Henry RJ. 2024. An improved haplotype resolved genome reveals more rice genes. Tropical Plants 3.

Aganezov S, Yan SM, Soto DC, Kirsche M, Zarate S, Avdeyev P, Taylor DJ, Shafin K, Shumate A, Xiao C. 2022. A complete reference genome improves analysis of human genetic variation. Science 376: eabl3533.

Altemose N, Logsdon GA, Bzikadze AV, Sidhwani P, Langley SA, Caldas GV, Hoyt SJ, Uralsky L, Ryabov FD, Shew CJ et al. 2022. Complete genomic and epigenetic maps of human centromeres. Science 376: eabl4178.

Arumuganathan K, Earle ED. 1991. Nuclear DNA content of some important plant species. Plant Molecular Biology Reporter 9: 208–218.

Bachmann K. 1972. Genome size in mammals. Chromosoma 37: 85–93.

Bao J, Zhang H, Wang F, Li L, Zhu X, Xu J, Wang Y, Liu Z, Zhai G, Xu H et al. 2024. Telomere-to-telomere genome assemblies of two Chinese Baijiu-brewing sorghum landraces. Plant Communications 10.1016/j.xplc.2024.100933: 100933.

Baranyi M, Greilhuber J, Swięcicki WK. 1996. Genome size in wild Pisum species. Theoretical and Applied Genetics 93-93: 717-721.

Bennett MD. 2003. Comparisons with Caenorhabditis (100 Mb) and Drosophila (175 Mb) Using Flow Cytometry Show Genome Size in Arabidopsis to be 157 Mb and thus 25 % Larger than the Arabidopsis Genome Initiative Estimate of 125 Mb. Annals of Botany 91: 547–557.

Bennett MD, Smith J. 1991. Nuclear DNA amounts in angiosperms. Philosophical Transactions: Biological Sciences: 309–345.

Bennett MD, Smith JB. 1976. Nuclear DNA amounts in angiosperms. Philosophical Transactions of the Royal Society of London B, Biological Sciences 274: 227–274.

Chang X, He X, Li J, Liu Z, Pi R, Luo X, Wang R, Hu X, Lu S, Zhang X et al. 2024. High-quality Gossypium hirsutum and Gossypium barbadense genome assemblies reveal the landscape and evolution of centromeres. Plant Communications 5: 100722.

Chao K-H, Zimin AV, Pertea M, Salzberg SL. 2023a. The first gapless, reference-quality, fully annotated genome from a Southern Han Chinese individual. G3 Genes|Genomes|Genetics 13.

Chao K-H, Zimin AV, Pertea M, Salzberg SL. 2023b. The first gapless, reference-quality, fully annotated genome from a Southern Han Chinese individual. G3: Genes, Genomes, Genetics 13: jkac321.

Chen J, Wang Z, Tan K, Huang W, Shi J, Li T, Hu J, Wang K, Wang C, Xin B et al. 2023. A complete telomere-to-telomere assembly of the maize genome. Nature Genetics 55: 1221–1231.

Cuzzoni E, Ferretti L, Giordani C, Castiglione S, Sala F. 1990. A repeated chromosomal DNA sequence is amplified as a circular extrachromosomal molecule in rice (Oryza sativa L.). Molecular and General Genetics MGG 222: 58–64.

Deschamps S, Zhang Y, Llaca V, Ye L, Sanyal A, King M, May G, Lin H. 2018. A chromosome-scale assembly of the sorghum genome using nanopore sequencing and optical mapping. Nature Communications 9: 4844.

Ding Y, Wang Y, Xu J, Jiang F, Li W, Zhang Q, Yang L, Zhao Z, Cheng B, Cao N. 2024. A telomere-to-telomere genome assembly of Hongyingzi, a sorghum cultivar used for Chinese Baijiu production. The Crop Journal 12: 635–640.

Doležel J, Bartoš J. 2005. Plant DNA Flow Cytometry and Estimation of Nuclear Genome Size. Annals of Botany 95: 99–110.

Doležel J, Bartoš J, Voglmayr H, Greilhuber J. 2003. Nuclear DNA content and genome size of trout and human. Cytometry Part A 51: 127–128.

Doležel J, Greilhuber J. 2010. Nuclear genome size: Are we getting closer? Cytometry Part A 77A: 635–642.

Doležel J, Greilhuber J, Lucretti S, Meister A, Lysák M, Nardi L, Obermayer R. 1998. Plant genome size estimation by flow cytometry: inter-laboratory comparison. Annals of botany 82: 17–26.

Dolezel J, Greilhuber J, Suda J. 2007. Flow cytometry with plant cells: analysis of genes, chromosomes and genomes. John Wiley & Sons.

Doležel J, Greilhuber J, Suda J. 2007. Estimation of nuclear DNA content in plants using flow cytometry. Nature Protocols 2: 2233–2244.

Doležel J, Sgorbati S, Lucretti S. 1992. Comparison of three DNA fluorochromes for flow cytometric estimation of nuclear DNA content in plants. Physiologia Plantarum 85: 625–631.

Galbraith DW, Harkins KR, Knapp S. 1991. Systemic endopolyploidy in Arabidopsis thaliana. Plant Physiology 96: 985–989.

Galbraith DW, Harkins KR, Maddox JM, Ayres NM, Sharma DP, Firoozabady E. 1983. Rapid flow cytometric analysis of the cell cycle in intact plant tissues. Science 220: 1049–1051.

Gladman N, Goodwin S, Chougule K, McCombie WR, Ware D. 2023. Era of gapless plant genomes: Innovations in sequencing and mapping technologies revolutionize genomics and breeding. Current Opinion in Biotechnology 79: 102886.

Goff SA, Ricke D, Lan T-H, Presting G, Wang R, Dunn M, Glazebrook J, Sessions A, Oeller P, Varma H. 2002. A draft sequence of the rice genome (Oryza sativa L. ssp. japonica). Science 296: 92–100.

Guo L, Chen W, Yan M, Chen S, Sun J, Wang J, Meng D, Li J, Zhang L. 2024. The complete genome assembly of Nicotiana benthamiana reveals genetic and epigenetic landscape of centromeres.

Hendrix B, Stewart JM. 2005. Estimation of the Nuclear DNA Content of Gossypium Species. Annals of Botany 95: 789–797.

Hou X, Wang D, Cheng Z, Wang Y, Jiao Y. 2022. A near-complete assembly of an Arabidopsis thaliana genome. Molecular Plant 15: 1247–1250.

Hu Y, Chen J, Fang L, Zhang Z, Ma W, Niu Y, Ju L, Deng J, Zhao T, Lian J. 2019. Gossypium barbadense and Gossypium hirsutum genomes provide insights into the origin and evolution of allotetraploid cotton. Nature genetics 51: 739–748.

Huang G, Wu Z, Percy RG, Bai M, Li Y, Frelichowski JE, Hu J, Wang K, Yu JZ, Zhu Y. 2020. Genome sequence of Gossypium herbaceum and genome updates of Gossypium arboreum and Gossypium hirsutum provide insights into cotton A-genome evolution. Nature genetics 52: 516–524.

Hussain Z, Sun Y, Shah S, Khan H, Ali S, Iqbal A, Zia M, Ali S. 2021. The dynamics of genome size and GC contents evolution in genus Nicotiana. Brazilian Journal of Biology 83: e245372.

Ihaka R, Gentleman R. 1996. R: a language for data analysis and graphics. Journal of computational and graphical statistics 5: 299–314.

Iyengar GAS, Sen SK. 1978. Nuclear DNA content of several wild and cultivated Oryza species. Environmental and Experimental Botany 18: 219–224.

Jarvis ED, Formenti G, Rhie A, Guarracino A, Yang C, Wood J, Tracey A, Thibaud-Nissen F, Vollger MR, Porubsky D. 2022. Semi-automated assembly of high-quality diploid human reference genomes. Nature 611: 519–531.

Johnston JS, Bennett MD, Rayburn AL, Galbraith DW, Price HJ. 1999. Reference standards for determination of DNA content of plant nuclei. American Journal of Botany 86: 609–613.

Jordan GJ, Carpenter RJ, Koutoulis A, Price A, Brodribb TJ. 2015. Environmental adaptation in stomatal size independent of the effects of genome size. New Phytologist 205: 608–617.

Kang M, Wu H, Liu H, Liu W, Zhu M, Han Y, Liu W, Chen C, Song Y, Tan L et al. 2023. The pan-genome and local adaptation of Arabidopsis thaliana. Nature Communications 14: 6259.

Kinoshita Y, Ohnishi N, Yamada Y, Kunisada T, Yamagishi H. 1985. Extrachromosomal Circular DNA from Nuclear Fraction of Higher Plants. Plant and Cell Physiology 26: 1401–1409.

Ko S-R, Lee S, Koo H, Seo H, Yu J, Kim Y-M, Kwon S-Y, Shin A-Y. 2024. High-quality chromosome-level genome assembly of Nicotiana benthamiana. Scientific Data 11: 386.

Koutecký P, Smith T, Loureiro J, Kron P. 2023. Best practices for instrument settings and raw data analysis in plant flow cytometry. Cytometry Part A.

Kreplak J, Madoui M-A, Cápal P, Novák P, Labadie K, Aubert G, Bayer PE, Gali KK, Syme RA, Main D. 2019. A reference genome for pea provides insight into legume genome evolution. Nature genetics 51: 1411–1422.

Krisai R, Greilhuber J. 1997. Cochlearia pyrenaica DC., das Löffelkraut, in Oberösterreich:(mit Anmerkungen zur Karyologie und zur Genomgröße). Biologiezentrum d. Oberösterr. Landesmuseums.

Kurotani K-i, Hirakawa H, Shirasawa K, Tanizawa Y, Nakamura Y, Isobe S, Notaguchi M. 2023. Genome Sequence and Analysis of Nicotiana benthamiana, the Model Plant for Interactions between Organisms. Plant and Cell Physiology 64: 248–257.

Lander ES, Linton LM, Birren B, Nusbaum C, Zody MC, Baldwin J, Devon K, Dewar K, Doyle M, FitzHugh W et al. 2001. Initial sequencing and analysis of the human genome. Nature 409: 860–921.

Laurie DA, Bennett MD. 1985. Nuclear DNA content in the genera Zea and Sorghum. Intergeneric, interspecific and intraspecific variation. Heredity 55: 307–313.

Leitch IJ JE, Pelicer J, Hidalgo O, Bennett MD. 2019. Angiosperm DNA C-values database (release 9.0, Apr 2019). 10.1111/nph.16261.

Li F, Fan G, Lu C, Xiao G, Zou C, Kohel RJ, Ma Z, Shang H, Ma X, Wu J et al. 2015. Genome sequence of cultivated Upland cotton (Gossypium hirsutum TM-1) provides insights into genome evolution. Nature Biotechnology 33: 524–530.

Li H, Durbin R. 2024. Genome assembly in the telomere-to-telomere era. Nature Reviews Genetics doi:10.1038/s41576-024-00718-w.

Lian Q, Huettel B, Walkemeier B, Mayjonade B, Lopez-Roques C, Gil L, Roux F, Schneeberger K, Mercier R. 2024. A pan-genome of 69 Arabidopsis thaliana accessions reveals a conserved genome structure throughout the global species range. Nature Genetics 56: 982–991.

Loureiro J, Rodriguez E, Doležel J, Santos C. 2007. Two new nuclear isolation buffers for plant DNA flow cytometry: a test with 37 species. Annals of botany 100: 875–888.

Lyndon RF. 1963. Changes in the Nucleus during Cellular Development in the Pea Seedling. Journal of Experimental Botany 14: 419–430.

Mccormick RF, Truong SK, Sreedasyam A, Jenkins J, Shu S, Sims D, Kennedy M, Amirebrahimi M, Weers BD, Mckinley B et al. 2018. The Sorghum bicolor reference genome: improved assembly, gene annotations, a transcriptome atlas, and signatures of genome organization. The Plant Journal 93: 338–354.

Michaelson MJ, Price HJ, Ellison JR, Johnston JS. 1991. COMPARISON OF PLANT DNA CONTENTS DETERMINED BY FEULGEN MICROSPECTROPHOTOMETRY AND LASER FLOW CYTOMETRY. American Journal of Botany 78: 183–188.

Miga KH, Koren S, Rhie A, Vollger MR, Gershman A, Bzikadze A, Brooks S, Howe E, Porubsky D, Logsdon GA et al. 2020. Telomere-to-telomere assembly of a complete human X chromosome. Nature 585: 79–84.

Mo C, Wang H, Wei M, Zeng Q, Zhang X, Fei Z, Zhang Y, Kong Q. 2024. Complete genome assembly provides a high-quality skeleton for pan-NLRome construction in melon. The Plant Journal 118: 2249–2268.

Morales J, Pujar S, Loveland JE, Astashyn A, Bennett R, Berry A, Cox E, Davidson C, Ermolaeva O, Farrell CM et al. 2022. A joint NCBI and EMBL-EBI transcript set for clinical genomics and research. Nature 604: 310–315.

Narayan R. 1987. Nuclear DNA changes, genome differentiation and evolution in Nicotiana (Solanaceae). Plant Systematics and Evolution 157: 161–180.

NCBI. 2024. Assembly [Internet]. Bethesda (MD): National Library of Medicine (US), National Centre of Biotechnology Information; [1988] - Assembly GCA_028009825.2, Genome assembly Col-CC [cited 2024/09/02]. Available from https://www.ncbi.nlm.nih.gov/datasets/genome/GCA_028009825.2/.

Nurk S, Koren S, Rhie A, Rautiainen M, Bzikadze AV, Mikheenko A, Vollger MR, Altemose N, Uralsky L, Gershman A et al. 2022. The complete sequence of a human genome. Science 376: 44–53.

Paterson AH, Bowers JE, Bruggmann R, Dubchak I, Grimwood J, Gundlach H, Haberer G, Hellsten U, Mitros T, Poliakov A et al. 2009. The Sorghum bicolor genome and the diversification of grasses. Nature 457: 551–556.

Praça-Fontes MM, Carvalho CR, Clarindo WR, Cruz CD. 2011. Revisiting the DNA C-values of the genome size-standards used in plant flow cytometry to choose the “best primary standards”. Plant cell reports 30: 1183–1191.

Ranawaka B, An J, Lorenc MT, Jung H, Sulli M, Aprea G, Roden S, Llaca V, Hayashi S, Asadyar L et al. 2023. A multi-omic Nicotiana benthamiana resource for fundamental research and biotechnology. Nature Plants 9: 1558–1571.

Reiser L, Bakker E, Subramaniam S, Chen X, Sawant S, Khosa K, Prithvi T, Berardini TZ. 2024. The Arabidopsis Information Resource in 2024. Genetics 227.

Rhie A, Nurk S, Cechova M, Hoyt SJ, Taylor DJ, Altemose N, Hook PW, Koren S, Rautiainen M, Alexandrov IA et al. 2023. The complete sequence of a human Y chromosome. Nature 621: 344–354.

Sadhu A, Bhadra S, Bandyopadhyay M. 2016. Novel nuclei isolation buffer for flow cytometric genome size estimation of Zingiberaceae: a comparison with common isolation buffers. Annals of Botany 118: 1057–1070.

Sahu SK, Liu H. 2023. Long-read sequencing (method of the year 2022): The way forward for plant omics research. Molecular Plant 16: 791–793.

Sasaki T. 2005. The map-based sequence of the rice genome. Nature 436: 793–800.

Schneider VA, Graves-Lindsay T, Howe K, Bouk N, Chen H-C, Kitts PA, Murphy TD, Pruitt KD, Thibaud-Nissen F, Albracht D. 2017. Evaluation of GRCh38 and de novo haploid genome assemblies demonstrates the enduring quality of the reference assembly. Genome research 27: 849–864.

Shang L, He W, Wang T, Yang Y, Xu Q, Zhao X, Yang L, Zhang H, Li X, Lv Y. 2023. A complete assembly of the rice Nipponbare reference genome. Molecular Plant 16: 1232–1236.

Sliwinska E, Loureiro J, Leitch IJ, Šmarda P, Bainard J, Bureš P, Chumová Z, Horová L, Koutecký P, Lučanová M et al. 2022. ApplicationLJbased guidelines for best practices in plant flow cytometry. Cytometry Part A 101: 749–781.

Song J-M, Xie W-Z, Wang S, Guo Y-X, Koo D-H, Kudrna D, Gong C, Huang Y, Feng J-W, Zhang W et al. 2021. Two gap-free reference genomes and a global view of the centromere architecture in rice. Molecular Plant 14: 1757–1767.

Soni A, Constantin L, Furtado A, Henry R. 2024. A flow cytometry protocol for accurate and precise measurement of plant genome size using frozen material. bioRxiv: 2024.2002. 2014.580322.

Tao Y, Luo H, Xu J, Cruickshank A, Zhao X, Teng F, Hathorn A, Wu X, Liu Y, Shatte T. 2021. Extensive variation within the pan-genome of cultivated and wild sorghum. Nature Plants 7: 766–773.

Temsch EM, Koutecký P, Urfus T, Šmarda P, Doležel J. 2022. Reference standards for flow cytometric estimation of absolute nuclear content in plants. Cytometry Part A 101: 710–724.

The Arabidopsis Genome I. 2000. Analysis of the genome sequence of the flowering plant Arabidopsis thaliana. Nature 408: 796–815.

Tiersch TR, Chandler RW, Wachtel SS, Elias S. 1989. Reference standards for flow cytometry and application in comparative studies of nuclear DNA content. Cytometry 10: 706–710.

Van’T Hof J. 1965. Relationships between mitotic cycle duration, S period duration and the average rate of DNA synthesis in the root meristem cells of several plants. Experimental Cell Research 39: 48–58.

Vollger MR, Guitart X, Dishuck PC, Mercuri L, Harvey WT, Gershman A, Diekhans M, Sulovari A, Munson KM, Lewis AP et al. 2022. Segmental duplications and their variation in a complete human genome. Science 376: eabj6965.

Wang B, Jiao Y, Chougule K, Olson A, Huang J, Llaca V, Fengler K, Wei X, Wang L, Wang X et al. 2021. Pan-genome Analysis in Sorghum Highlights the Extent of Genomic Variation and Sugarcane Aphid Resistance Genes. doi:10.1101/2021.01.03.424980. Cold Spring Harbor Laboratory.

Wang J, Zhang Q, Tung J, Zhang X, Liu D, Deng Y, Tian Z, Chen H, Wang T, Yin W. 2024. High-quality assembled and annotated genomes of Nicotiana tabacum and Nicotiana benthamiana reveal chromosome evolution and changes in defense arsenals. Molecular Plant 17: 423–437.

Wang M, Tu L, Yuan D, Zhu D, Shen C, Li J, Liu F, Pei L, Wang P, Zhao G. 2019. Reference genome sequences of two cultivated allotetraploid cottons, Gossypium hirsutum and Gossypium barbadense. Nature genetics 51: 224–229.

Wickham H. 2011. ggplot2. Wiley interdisciplinary reviews: computational statistics 3: 180–185.

Xie L, Gong X, Yang K, Huang Y, Zhang S, Shen L, Sun Y, Wu D, Ye C, Zhu Q-H. 2024. Technology-enabled great leap in deciphering plant genomes. Nature Plants 10: 551–566.

Yang T, Liu R, Luo Y, Hu S, Wang D, Wang C, Pandey MK, Ge S, Xu Q, Li N. 2022. Improved pea reference genome and pan-genome highlight genomic features and evolutionary characteristics. Nature genetics 54: 1553–1563.

Zhang Y, Fu J, Wang K, Han X, Yan T, Su Y, Li Y, Lin Z, Qin P, Fu C et al. 2022. The telomere-to-telomere gap-free genome of four rice parents reveals SV and PAV patterns in hybrid rice breeding. Plant Biotechnology Journal 20: 1642–1644.

Zhuang J, Zhang Y, Zhou C, Fan D, Huang T, Feng Q, Lu Y, Zhao Y, Zhao Q, Han B. 2024. Dynamics of extrachromosomal circular DNA in rice. Nature Communications 15: 2413.

