## Supplemental material Figures S1-4 Tables S1-10 for "Re-calibration of flow cytometry standards for plant genome size estimation"

**Title**

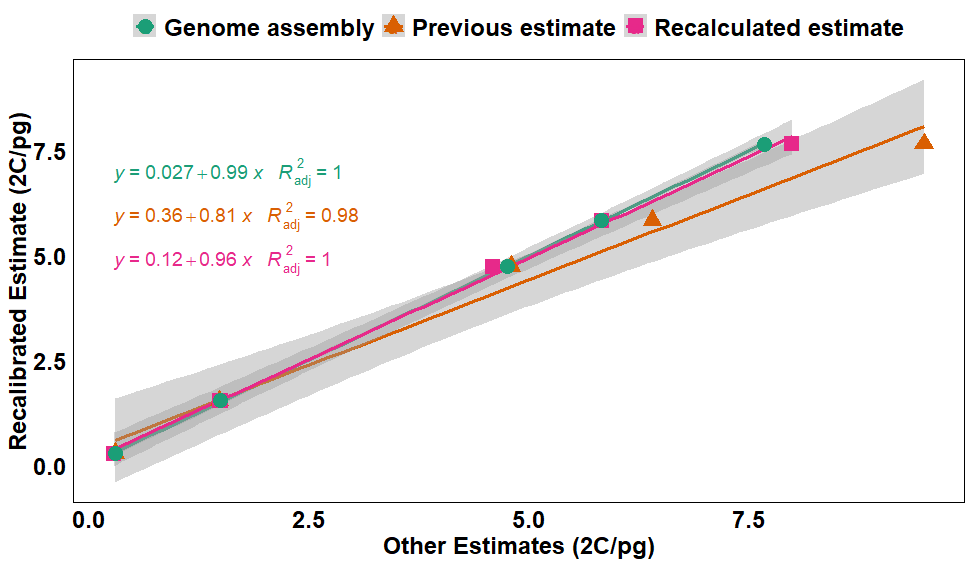

**Figure S1** Regression analysis of the recalibrated GS estimates against genome assembly-based estimates, previous estimates, and recalculated estimates for five plant reference standards. Higher slope values and smaller interecepts demonstrate strong linear relationships between the recalibrated estimate, recalculated estimates and genome assembly size.
